# Hair follicles modulate skin barrier function

**DOI:** 10.1101/2024.04.23.590728

**Authors:** Noah C. Ford, Rachel E. Benedeck, Matthew T. Mattoon, Jamie K. Peterson, Arlee L. Mesler, Natalia A. Veniaminova, Danielle J. Gardon, Shih-Ying Tsai, Yoshikazu Uchida, Sunny Y. Wong

## Abstract

Our skin provides a protective barrier that shields us from our environment. Barrier function is typically associated with interfollicular epidermis; however, whether hair follicles influence this process remains unclear. Here, we utilize a potent genetic tool to probe barrier function by conditionally ablating a quintessential epidermal barrier gene, *Abca12*, which is mutated in the most severe skin barrier disease, harlequin ichthyosis. With this tool, we deduced 4 ways by which hair follicles modulate skin barrier function. First, the upper hair follicle (uHF) forms a functioning barrier. Second, barrier disruption in the uHF elicits non-cell autonomous responses in the epidermis. Third, deleting *Abca12* in the uHF impairs desquamation and blocks sebum release. Finally, barrier perturbation causes uHF cells to move into the epidermis. Neutralizing Il17a, whose expression is enriched in the uHF, partially alleviated some disease phenotypes. Altogether, our findings implicate hair follicles as multi-faceted regulators of skin barrier function.

## Introduction

Our skin serves as a protective barrier that surrounds our internal organs while providing an interface for gas and liquid exchange. This barrier is formed when keratinocytes differentiate, stratify and undergo cornification, a specialized form of cell death that gives rise to corneocytes^1,2^. In the outermost layer of the skin, known as the stratum corneum, these corneocytes form a scaffold with other cross-linked structural proteins (“bricks”) and are surrounded by an amalgam of lipid-dominant lamellar sheets (“mortar”)^3^. Together, these “bricks and mortar” comprise the hydrophobic, insoluble barrier that is essential for life. With >100 inherited skin diseases linked to mutations in genes associated with barrier function, even subtle defects in this complicated process can have profound consequences^4^.

Ceramides are essential barrier lipids that constitute up to 50% of the total lipids by weight in the cornified envelope^5^. These sphingolipids are synthesized by keratinocytes and transported by membrane-enclosed organelles known as lamellar granules (LGs)^6^. In granular layer keratinocytes, LGs travel from the Golgi apparatus to the apical plasma membrane, where they fuse and release their contents into the interstitial spaces of the overlying stratum corneum. In addition to lipids, LGs deliver a variety of cargo, including an assortment of lipid- and protein-processing enzymes, proteases for desquamation, structural proteins and anti-microbial peptides^7^. As such, LGs serve as the main secretory organelles in the skin.

Inherited ichthyoses encompass a large class of genetic disorders rooted in defective cornification^8^. Common features include dysregulated barrier function, inflammation and hyperkeratosis. Among all skin barrier diseases, harlequin ichthyosis (HI) is the most severe and is caused by mutations in *ABCA12*, which encodes an ATP-driven lipid transporter that localizes to LGs^9,10^. In the absence of ABCA12, LGs fail to deliver lipids and desquamation enzymes to the stratum corneum^11,12^. Consequently, HI patients possess scaly, plate-like skin that is paradoxically thickened but dysfunctional, lacking essential barrier lipids. Although neonatal intensive care and retinoid therapy can partially relieve symptoms, HI patients suffer from debilitating lifelong disfiguration and increased susceptibility to infection and dehydration^13,14^.

Most studies on skin barrier function and disease have focused on the interfollicular epidermis (IFE). However, the hair follicle epithelium is continuous with the IFE, and hair follicle openings constitute up to 10% of the skin’s total surface area at body sites such as the face^15^. This upper hair follicle (uHF) domain, also known as the hair canal or infundibulum, is colonized by commensal microbes and secretes a variety of cytokines and chemokines to recruit immune cells^16–20^. The uHF also provides a passageway for sebaceous gland-derived lipids to leave the follicle and enter the skin’s surface, where these oils can also modulate barrier function^21,22^. Thus, given its unique anatomic location at the interface between the outer epidermis and the deeper hair follicle, the uHF likely serves as an active site of commerce in the skin.

Pharmacologically, the hair follicle is thought to be an important penetration route for topically applied drugs^15,23,24^. Topical nanoparticles have also been reported to accumulate within the uHF^25^, yet the influence of this domain on skin barrier function remains unclear^26^. Here, we utilize a potent genetic strategy—targeted ablation of *Abca12*—to probe the consequences of disrupting a canonical epidermal barrier protein within the hair follicle.

## Results

### The hair follicle forms a functional barrier

The epidermis and hair follicle are maintained by basal stem cell progenitors that give rise to differentiated suprabasal cells (**Figure 1A**). Previously, we reported that the inner, differentiated cells of the uHF express Keratin (K) 79, a defining marker of this region^27^ (**Figure 1B-C**). To further compare the IFE and uHF compartments, we first assessed the localization of proteins associated with barrier function (Abca12, Rab11a)^28^, desquamation (kallikrein (Klk) 6 and 7)^29^, proteolysis (cathepsin D (Ctsd))^30^ and differentiation (K10) in mouse dorsal skin. Although differentiated cells in both compartments express all markers, we noted that Klk6, Klk7 and Ctsd are especially enriched in K79+ uHF cells (**Figure 1D**, **S1A**). In contrast, the ceramide precursor glucosylceramide (GlcCer) is abundant in the upper layers of the IFE but reduced in the uHF (**Figure 1D**). These findings indicate that barrier-associated proteins are found in both the IFE and uHF; however, these domains are also molecularly, and possibly functionally, distinct.

**Figure 1.**
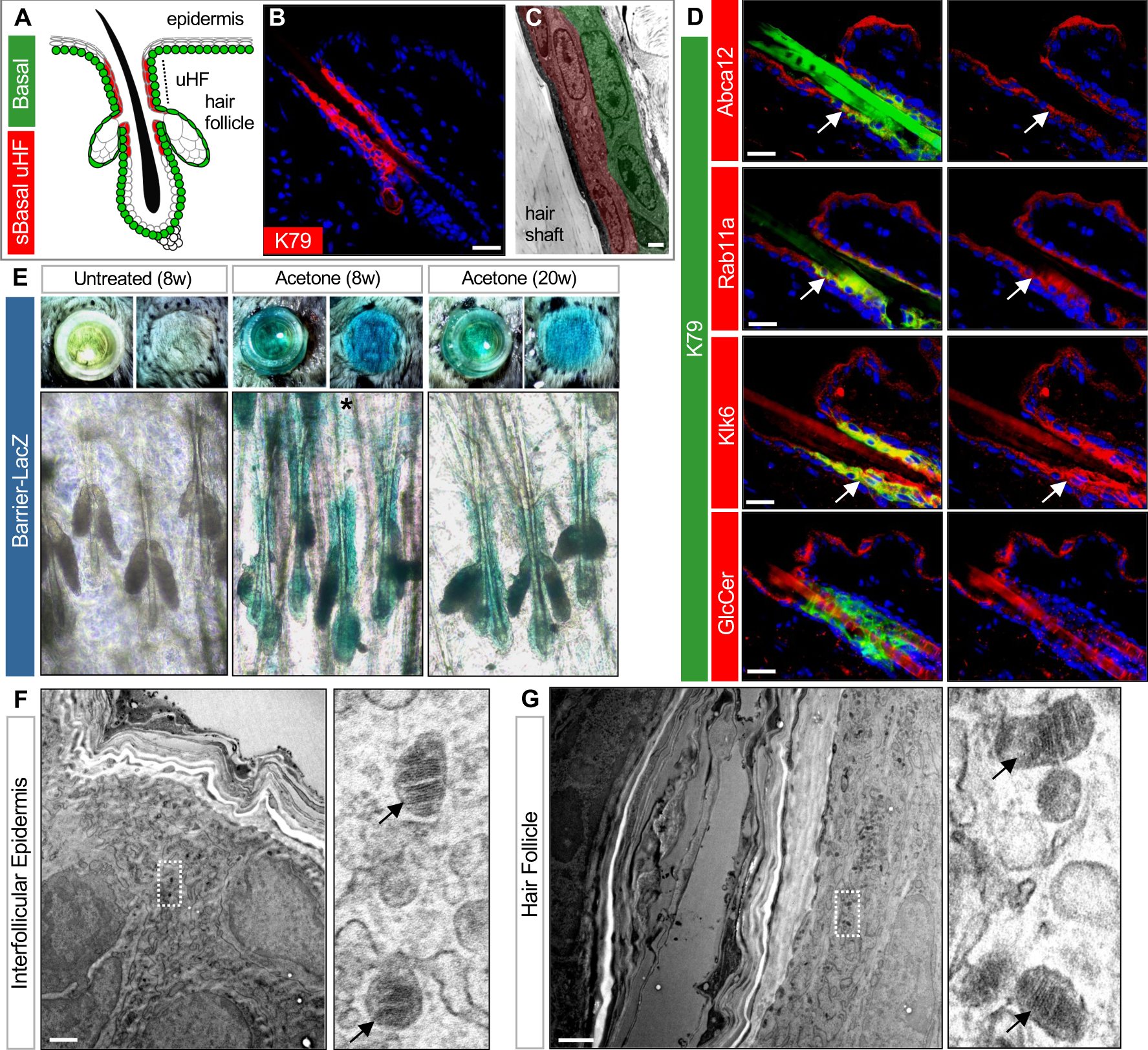
The hair follicle forms a functioning barrier. **A.** Schematic of telogen hair follicle. Green, basal progenitors. Red, K79+ suprabasal (sBasal) cells in the upper hair follicle (uHF). Note that K79 is also expressed by sebaceous glands and ducts^80^ (not highlighted here). **B.** Immunohistochemical (IHC) staining for K79 (red). **C.** Transmission electron microscopy (TEM) of uHF showing basal (green) and suprabasal (red) hair follicle epithelium and hair shaft. **D.** IHC for barrier-associated proteins (red) and K79 (green). Right panels show single-channel views of barrier-associated protein expression. Arrows, regions of overlap with K79+ uHF cells. **E.** Top panels, overhead views of skin following incubation with X-gal in cloning cylinders (left) or with cylinders removed (right). Bottom panels, whole mounts showing LacZ staining of untreated 8 week (w) old skin (left), acetone-treated 8w skin (middle), and acetone-treated 20w skin (right). Asterisk, occasional LacZ+ cells in the IFE. **F.** TEM of IFE. Right panel is a magnified view showing LGs (arrows). **G.** TEM of uHF. Right panel is a magnified view showing LGs (arrows). Scale bar for C, F, G, 1 μm. Scale bar for all others, 50 μm. See also **Figure S1**.

To visualize sites of barrier function, we next adapted an X-gal permeability assay originally described by Hardman et al.^31^ In this assay, barrier-competent skin excludes X-gal and remains unlabeled, whereas barrier-deficient skin turns blue from X-gal cleavage by endogenous β-galactosidase (LacZ). In 8-or 20-week old telogen mouse skin, we exposed the surface to X-gal loaded into cloning cylinders and observed no staining, indicating that the epidermis and hair follicle form a functional barrier (**Figure 1E**). However, when skin was pre-treated with acetone to perturb barrier lipids^32^ and then incubated with X-gal, extensive blue staining was seen in the hair follicle epithelium, with far less labeling in the epidermis, indicating selective barrier disruption in the follicle (**Figure 1E**).

Given these findings, we delved deeper into the ultrastructure of these domains, and observed granular cells containing LGs, as well as secreted lipid lamellae, in both the IFE and uHF (**Figure 1F-G**, **S1B-C**). Critically, whereas the IFE formed numerous cornified layers, these layers were less prominent towards the deeper follicle (**Figure S1D**). Altogether, these data suggest that while the hair follicle can enact a functioning barrier, the follicular barrier may be less robust than the epidermal barrier.

### Validating a genetic tool to disrupt barrier function

We next used a genetic strategy to probe barrier function. Since HI is the most severe skin barrier disease, we reasoned that targeted deletion of *Abca12*, the causative gene mutated in HI^9,10^, would provide us a tool to disrupt lamellar granules and interrogate barrier function in different epithelial sub-compartments. We therefore first generated mice harboring a null allele of *Abca12* that is constitutively disrupted by a *LacZ* insertion between exons 3 and 4, similar to previously described^33^ (**Figure S2A-B**). Newborn homozygous mutant pups (*Abca12-KO*) recapitulated HI features, including taut, shiny skin; thickened and compacted stratum corneum; and death from rapid dehydration^34–36^ (**Figure 2A**). We further confirmed loss of Abca12 by immunohistochemistry (IHC), and validated by X-Gal staining that the *Abca12* promoter is active in both newborn epidermis and developing hair canals (**Figure S2C**).

**Figure 2.**
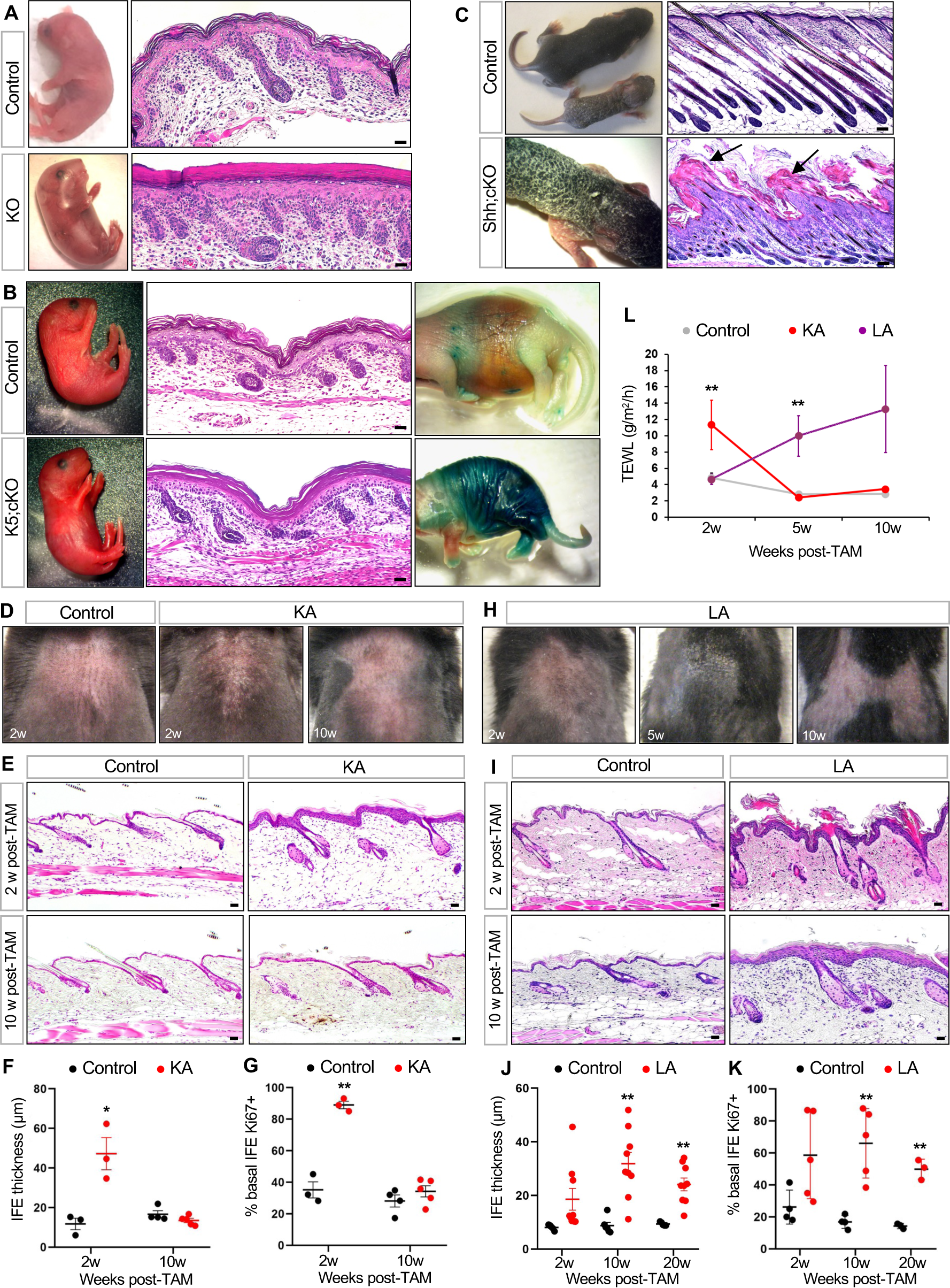
*Abca12* mutant mice provide a genetic tool to probe barrier function. **A.** Gross images of (left) and skin histology from (right) newborn *Abca12* KO pup or littermate control. **B.** Similar to (A), but comparing newborn *K5;cKO* pup and littermate control. Right-most panels, LacZ barrier assay, with blue staining indicating dysfunctional skin barrier^31^. Note that the cKO allele does not contain the *LacZ* transgene. **C.** Similar to (A), but comparing *Shh;cKO* pup and littermate control, on postnatal (P) day 7-8. **D.** Gross images of shaved dorsal skin from control or KA mice at the indicated number of weeks (w) post-TAM. **E.** Skin histology of control or KA mice at 2 weeks (top) or 10 weeks (bottom) post-TAM. **F.** Quantitation of IFE thickness in control (black) or KA (red) mice. **G.** Quantitation of basal IFE proliferation in control or KA mice. **H.** Gross images of shaved dorsal skin from LA mice, at the indicated times post-TAM. See **Figure S3B** for image of littermate control mouse. **I.** Skin histology of control or LA mice at 2 weeks (top) or 10 weeks (bottom) post-TAM. **J.** Quantitation of IFE thickness in control (black) or LA (red) mice. **K.** Quantitation of basal IFE proliferation in control or LA mice. **L.** TEWL measurements from control (gray), KA (red) and LA (purple) mice. For **F, G, J, K**: *, p < 0.05; **, p < 0.01 by unpaired t-test, comparing only samples from the same timepoint. n ≥ 3 mice per genotype, per timepoint. For **L**: **, p < 0.01 by one-way ANOVA with post hoc Tukey test, comparing only samples from the same timepoint. For each timepoint, n = 4-5 KA mice and 6-9 LA mice, and ≥ 14 control mice. Scale bar for C, 100 μm. Scale bar for all others, 50 μm. See also **Figure S2-S3**.

Since early lethality precluded further studies with KO mice, we next generated a conditional allele of *Abca12* (cKO) (**Figure S2A**). When homozygous *Abca12* cKO alleles were coupled with *Keratin 5* (K5) promoter-driven Cre recombinase to delete *Abca12* throughout the skin^37^, these *K5;cKO* pups also exhibited severe barrier defects, validating this floxed deletion allele (**Figure 2B**, **S2D**).

Finally, we coupled homozygous *Abca12* cKO alleles with *Sonic hedgehog* (Shh) promoter-driven Cre recombinase, which targets genetic deletion to developing hair follicle progenitors, but not to the epidermis^38^. In striking contrast to *Abca12-KO* and *K5;cKO* mice, newborn *Shh;cKO* pups were indistinguishable from littermate controls up until postnatal (P) day 4. Subsequently, these mutants became runted and developed thick, scaly skin (**Figure 2C**). Histological analysis revealed that while follicular downgrowth was largely unaffected, hair follicle openings were plugged with keratotic material (**Figure 2C**). Since *Shh;cKO* mutants failed to thrive beyond ∼P8, we grafted newborn mutant skin onto nude mice, and found that hair protrusion was impaired even after several weeks (**Figure S2E**). These findings confirm that the *Abca12* cKO allele can serve as a powerful genetic tool to disrupt at least 2 key features of normal skin—barrier function and desquamation.

### Divergent outcomes after deleting *Abca12* in different stem cell compartments

To probe the effects of deleting *Abca12* in adult skin, we generated 2 additional tamoxifen (TAM)-inducible mouse strains. We and others have previously shown that *K14* promoter-driven Cre^ERT^ causes recombination primarily in IFE stem cells, and less so in the follicle^39,40^ (**Figure S3A**). In contrast, *Lrig1* promoter-driven Cre^ERT^^2^ induces recombination in Lrig1+ stem cells that maintain the uHF^41,42^ (**Figure S3A**). We therefore generated *<u>K</u>14;<u>A</u>bca12-cKO* (KA) and *<u>L</u>rig1;<u>A</u>bca12-cKO* (LA) mice to compare the effects of deleting *Abca12* in the IFE and uHF, respectively, in 8-week old telogen skin.

In KA mice, we noticed the appearance of dry, flaky skin at 2 weeks post-TAM (**Figure 2D**). Unexpectedly, this phenotype disappeared by 10 weeks post-TAM (**Figure 2D**). These observations were confirmed histologically, which revealed that both epidermal thickness and proliferation increased sharply at 2 weeks post-TAM, but then returned to normal by 10 weeks post-TAM (**Figure 2E-G**).

In LA mice, we also observed dry, flaky skin starting at 2-5 weeks post-TAM (**Figure 2H**, **S3B**). Unlike KA mice, LA mutants displayed severe follicular hyperkeratosis, similar to *Shh;cKO* pups (**Figure 2I**). Although the overall appearance of some LA mice seemed to improve over time, all mutants retained chronically inflamed, reddish skin, and IHC revealed that the epidermis was hyperplastic and hyperproliferative even 20 weeks post-TAM (**Figure 2J-K**).

These trends—an immediate but reversible ichthyotic phenotype in KA mice, versus longer-lasting effects in LA mutants—coincided with rates of transepidermal water loss (TEWL), a measure of inside-out skin barrier function. Whereas KA mice exhibited a temporary spike in TEWL, LA mutants displayed a more gradual sustained increase (**Figure 2L**). These findings demonstrate that deleting *Abca12* in distinct IFE and hair follicle stem cell compartments causes divergent long-term outcomes in the skin. We explore the molecular and cellular basis for these differences below.

### Disrupting the hair follicle barrier induces non-cell autonomous responses

As mentioned above, the *Abca12* KO allele contains a *LacZ* insertion that simultaneously disrupts gene function while reporting on endogenous *Abca12* promoter activity (**Figure S2A**). In mice harboring both the KO and cKO alleles of *Abca12*—where one allele is constitutively inactivated by *<u>L</u>acZ* and the other is poised for deletion—targeted disruption of *Abca12* in the adult IFE using K14-Cre^ERT^ (KA<u>L</u> mice) caused a transient increase in *Abca12* promoter activity in the hyperplastic epidermis, which later waned as the skin returned to normal (**Figure 3A**). Unexpectedly, when *Abca12* was deleted in the uHF using Lrig1-Cre^ERT^^2^ (LA<u>L</u> mice), this also caused increased *Abca12* promoter activity in the IFE (**Figure 3A**). Notably, this effect was longer-lasting, coincided with sustained epidermal hyperplasia, and occurred outside of the uHF, where *Abca12* was initially deleted.

**Figure 3.**
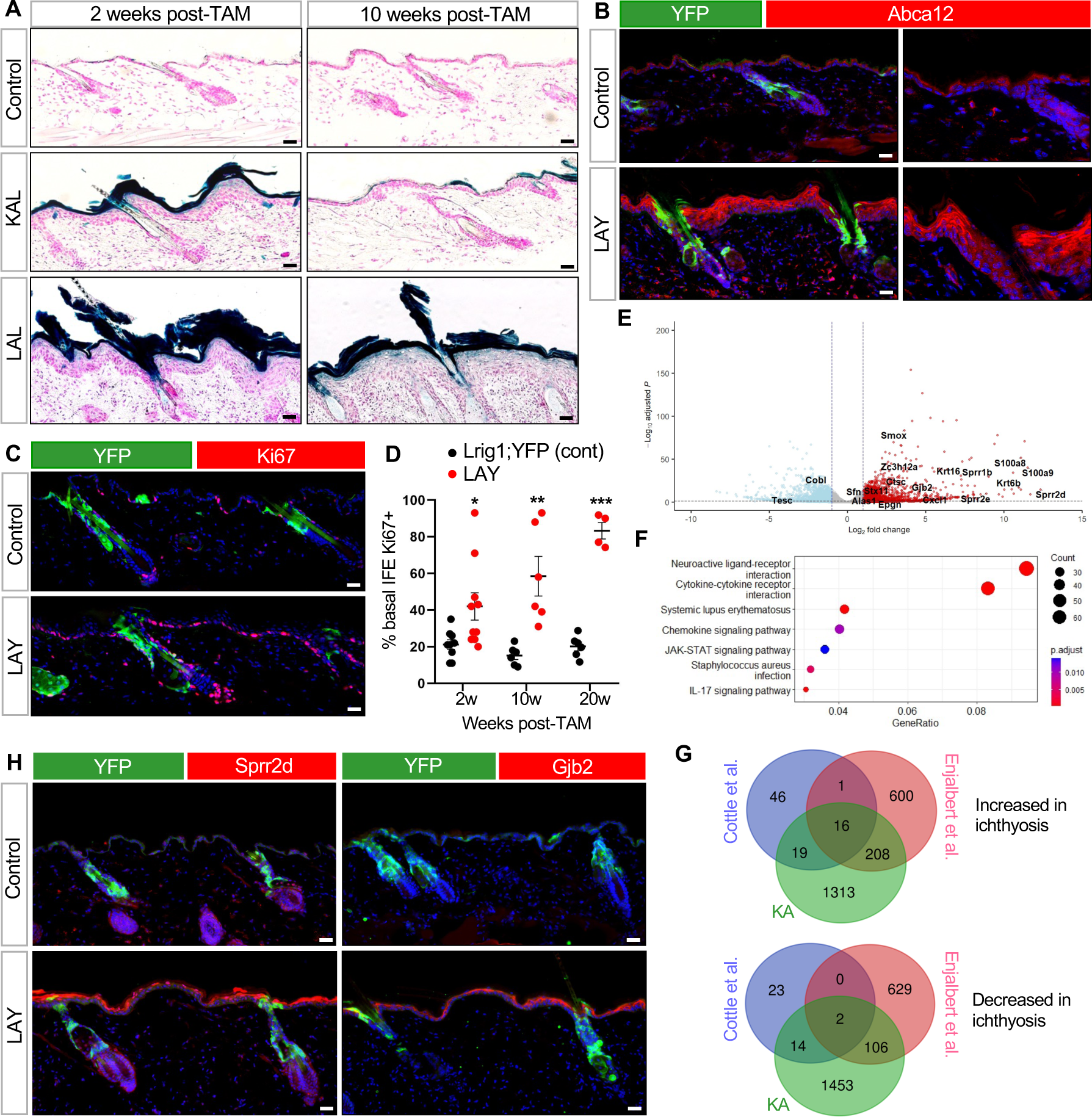
*Abca12* deletion in the uHF induces non-cell autonomous responses in the IFE. **A.** LacZ staining, as a read-out for *Abca12* promoter activity, in *K14-Cre^ERT^;Abca12-KO/cKO* (KAL) or *Lrig1-Cre^ERT^*^2^*;Abca12-KO/cKO* (LAL) mice. Controls are *Abca12-KO/cKO* mice lacking Cre. **B.** IHC for Abca12 in *Lrig1-Cre^ERT^*^2^*;Abca12-c/c;ROSA-YFP* (LAY) mice, 2 weeks post-TAM. Control mice are similar to LAY mutants, but possess one wild-type copy of *Abca12* (*Abca12-c/+*). Right panels are magnified, single-channel views showing non-cell autonomous upregulation of Abca12 (red) in LAY skin that does not overlap with YFP+ uHF cells. **C.** Same as (B) but with IHC for Ki67 (red), 10 weeks post-TAM. **D.** Quantitation of proliferation in YFP-negative, basal IFE cells. **E.** Volcano plot showing upregulated (red) and downregulated (blue) DEGs in KA mice and control *Abca12-c/c* mice lacking Cre. **F.** KEGG enrichment analysis of all DEGs. **G.** Venn diagrams showing overlap of DEGs in KA mice, HI patients^44^ and *Abca12*-mutant embryonic mice^43^. Overlapping genes from the 3 data-sets are highlighted in (E). **H.** Same as (B) but with IHC for Sprr2d (red, left panels) or Gjb2 (red, right panels), 2 weeks post-TAM. For **D**: *, p < 0.05; **, p < 0.01; ***, p < 0.001 by unpaired t-test, comparing only samples from the same timepoint. n ≥ 4 mice per genotype, per timepoint. Scale bar, 50 μm. See also **Figure S3**.

We next confirmed these observations using *Lrig1-Cre^ERT^*^2^ mice harboring homozygous *Abca12* cKO alleles and a *<u>Y</u>FP* reporter to identify cells that had undergone Cre-mediated recombination (LA<u>Y</u> mice). Here, IHC staining confirmed that Abca12 protein and cell proliferation were both increased in non-recombined, YFP-negative cells in the IFE, suggesting a non-cell autonomous effect (**Figure 3B-D**).

To broaden our scope, we performed bulk RNA-sequencing (RNA-seq) to assess transcriptional changes in KA mice at 2 weeks post-TAM. This identified 1,556 upregulated genes, including numerous genes involved with epidermal differentiation, barrier function and wound healing (**Figure 3E**). Pathway analysis further revealed significant enrichment for signaling modules associated with lupus erythematosus, JAK-STAT and IL17 (**Figure 3F**). Finally, we overlapped our RNA-seq dataset with those previously generated by Cottle et al., from *Abca12*-mutant mouse embryos^43^ and by Enjalbert et al., from HI patients^44^. This identified 16 common upregulated genes, including *Sprr2d*, *S100a9* and *Gjb2* (Connexin 26), which is mutated in keratitis-ichthyosis-deafness (KID) syndrome^45^ (**Figure 3E, G**).

To determine whether these genes can also be induced non-cell autonomously, we performed IHC for Sprr2d, S100a9 and Gjb2 in LAY mice. Indeed, when *Abca12* was deleted in the uHF, all 3 proteins became increased in non-recombined cells in the IFE (**Figure 3H, S3C**). These findings demonstrate that disrupting the follicular barrier causes the IFE to upregulate proteins involved with barrier function, possibly as a compensatory way to restore overall skin homeostasis.

### Altered sebum secretion and hair loss after deleting *Abca12* in the uHF

Our previous studies showed that disrupting epidermal differentiation indirectly causes enlargement of sebaceous glands (SGs), which are hair follicle-associated appendages that release oily secretions known as sebum into the uHF^39^. We therefore examined whether *Abca12* mutants also possess expanded SGs, and indeed found that SGs were enlarged in LA mice (**Figure 4A-B**). These effects are likely indirect, since deleting *Abca12* in the IFE also expanded SGs in KA mice, consistent with our previously published findings (**Figure 4C-D**).

**Figure 4.**
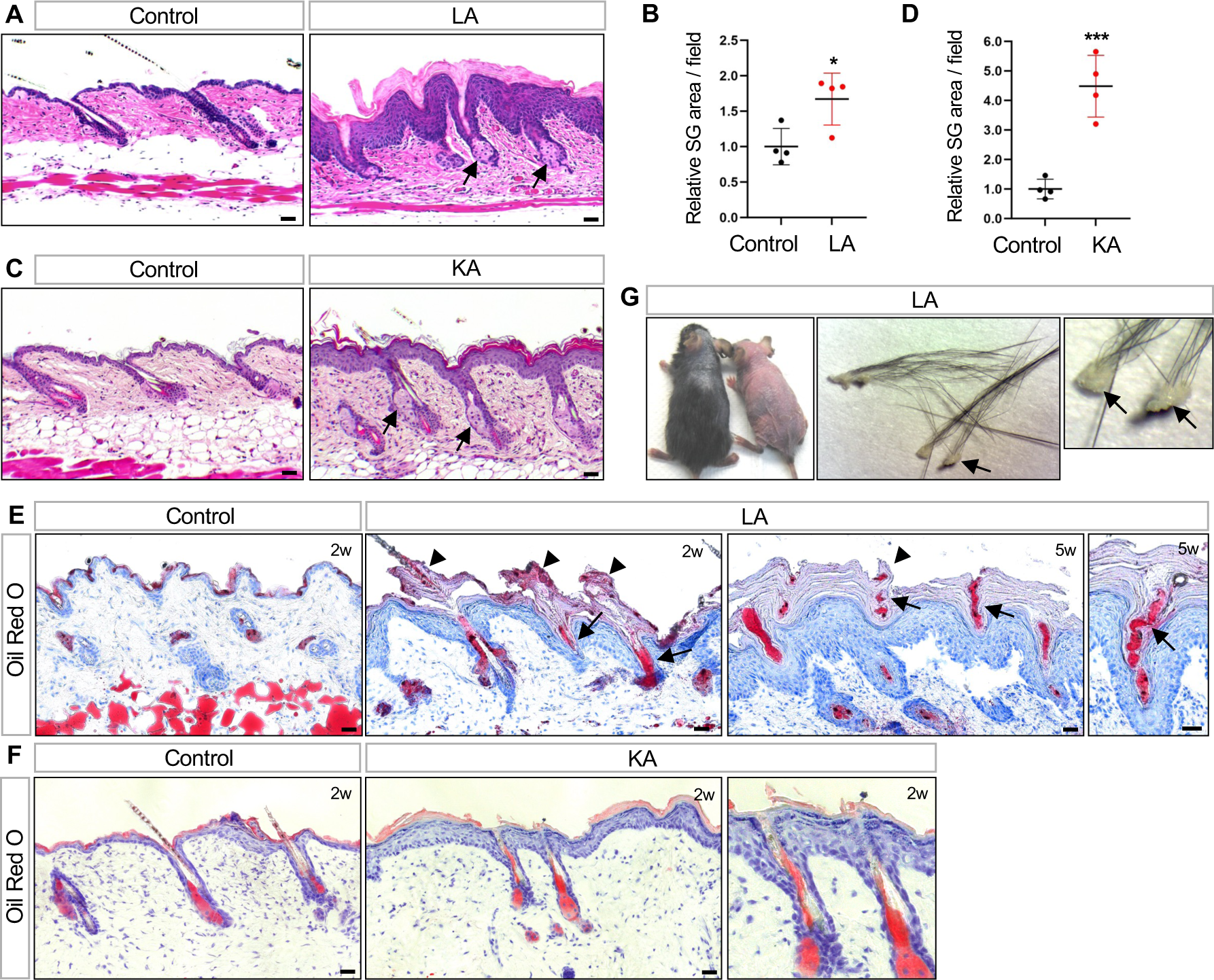
Impaired sebum secretion and hair loss following *Abca12* deletion in the uHF. **A.** Histology showing enlarged SGs (arrows) in LA skin. **B.** Quantitation of SG area in LA (red) or control (black) skin. **C.** Histology showing enlarged SGs (arrows) in KA skin. **D.** Quantitation of SG area in KA (red) or control (black) skin. **E.** Oil red O staining showing sebum lipids (arrows) trapped by hyperkeratotic material (arrowheads) in LA mice, 2-5 weeks (w) post-TAM. **F.** Same as (E) but for KA mice, 2 weeks post-TAM. Note that follicular hyperkeratosis and sebum occlusion are not observed even though the adjacent IFE is hyperplastic. Right panel is magnified view of the center. **G.** Photo showing hair loss in two LA mutant mice, both 5 weeks post-TAM (left photo). Enlarged views of hair clumps with oily plugs (arrows) at the proximal end (middle and right photos). For **B**, **D**: *, p < 0.05; ***, p < 0.001 by unpaired t-test. n = 4 mice per genotype. Scale bar, 50 μm.

To monitor lipid output from these enlarged SGs, we next stained skin sections with the lipophilic dye Oil Red O. Notably, LA skin exhibited extensive follicular hyperkeratosis that blocked the outflow of sebum and caused entrapment of oils within the uHF (**Figure 4E**). In contrast, these phenotypes were rarely seen in KA skin, even though the IFE was hyperplastic (**Figure 4F**). These results indicate that the uHF specifically must undergo constant desquamation to allow unimpeded sebum release. When this process is blocked, a subset of LA and LAL mice also developed hair loss, with shed hairs retaining oily keratotic plugs, suggesting that fracturing had occurred near sites of occlusion in the uHF (**Figure 4G**).

### Hair follicle-derived mutant cells enter the epidermis

Thus far, we have shown that hair follicles form a functioning barrier; modulate expression of barrier proteins in the IFE non-cell autonomously; and control the release of sebum, which also affects the skin’s barrier. However, a lingering question still remains: Why does deleting *Abca12* in the IFE cause transient ichthyotic phenotypes, whereas deleting *Abca12* in the uHF cause longer-lasting effects?

To answer this, we traced the fate of mutant (*Abca12-c/c*) or control (*Abca12-c/+*) cells over time, again using the Cre-inducible *<u>Y</u>FP* reporter. Following Cre-mediated recombination in the IFE (KA<u>Y</u> mice), we initially observed numerous YFP+ mutant cell clones in the epidermis after 2 weeks (**Figure 5A**, **S3D**). By 10 weeks post-TAM, however, the abundance of labeled mutant cells fell dramatically, coinciding with the phenotypic recovery seen in these animals (**Figure 5B**). In contrast, the abundance of YFP+ cells in control mice remained stable over time (**Figure 5A-B**).

**Figure 5.**
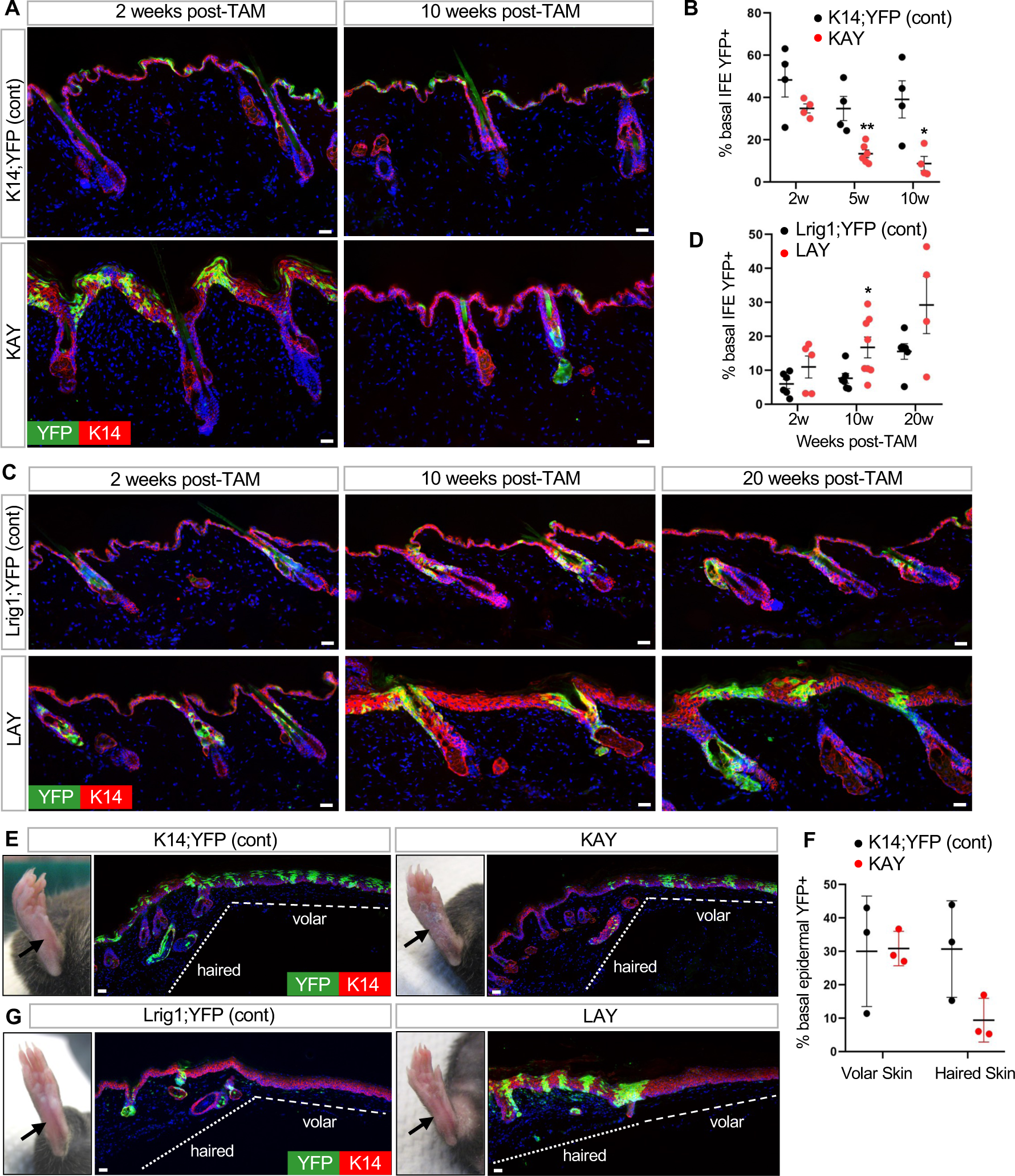
Mutant uHF cells move into the IFE. **A.** IHC to trace recombined YFP+ cells (green) in *K14-Cre^ERT^;Abca12-c/c;ROSA-YFP (KAY)* mice. Control mice are similar to KAY mutants, but possess one wild-type copy of *Abca12* (*Abca12-c/+*). **B.** Quantitation of basal YFP+ cells in the IFE of KAY (red) or control (black) mice. **C.** IHC to identify recombined YFP+ cells (green) in LAY mice or controls. Genotypes are identical to those described in Figure 3B. **D.** Quantitation of basal YFP+ cells in the IFE of LAY (red) or control (black) mice. **E.** IHC for YFP+ cells (green) in KAY or control paw volar skin, 15 weeks post-TAM. **F.** Quantitation of epidermal basal YFP+ cells in volar and adjacent haired skin, from KAY (red) or control (black) mice. n = 3 mice per genotype. **G.** IHC for YFP+ cells (green) in LAY or control volar skin, 10 weeks post-TAM. No recombination occurred in volar skin epidermis. For **B**, **D**: *, p < 0.05; **, p < 0.01 by unpaired t-test, comparing only samples from the same timepoint. n ≥ 4 mice per genotype, per timepoint. Scale bar, 50 μm. See also **Figure S3**.

A dramatically different story emerged when we traced the fate of *Abca12* mutant cells in the uHF (LA<u>Y</u> mice). Here, mutant cells not only persisted long-term in the uHF, but also spread into the IFE (**Figure 5C-D**, **S3E**). Labeled uHF cells in control mice also entered the IFE, but did so more gradually (**Figure 5C-D**).

These divergent outcomes—loss of mutant cells from the IFE, and expansion of mutant cells from the uHF—might be explained if Abca12 is required by epidermal cells to maintain competitive fitness. However, when we examined the fate of mutant cells in paw skin epidermis, a region devoid of hair follicles, these cells persisted up to 10 weeks post-TAM in KAY mice (**Figure 5E-G**). This indicates that loss of *Abca12*-deficient cells occurs only in haired epidermis, arguing against a general requirement for Abca12 to maintain cell fitness. Instead, these observations suggest that Abca12 may possess critical skin site-specific functions. In haired skin IFE, mutant cells may become out-competed by non-recombined neighboring cells or possibly diluted by cells originating from the uHF, as is seen after skin wounding^42,46^. In LAY mice, mutant cells persist in the uHF and spread into the IFE to maintain the ichthyotic phenotype (**Figure 5C, 5G**). The movement of these mutant cells across compartments thereby suggests yet another mechanism by which hair follicles can modulate skin barrier function.

### Disease features are partially driven by Il17a

We next sought to identify factors that drive the pleiotropic ichthyotic phenotypes seen in our models. As mentioned above, our RNA-seq and functional enrichment analysis revealed that Il17 signaling is increased in KA mice at 2 weeks post-TAM, when peak disease features are observed (**Figure 3F**). This signature was partially driven by heightened expression of *Il17a* (log2 fold-change of 8.24). By RNA-scope *in situ* staining, we confirmed that *Il17a* is elevated throughout the IFE in both KA and LA mutants (**Figure 6A**). Unexpectedly, in both models, we noticed that *Il17a* RNA is especially abundant in the uHF, and in SGs from LA mutants (**Figure 6A**). *Il17a* is also enriched in the uHF in non-ichthyotic control skin (**Figure 6A**).

**Figure 6.**
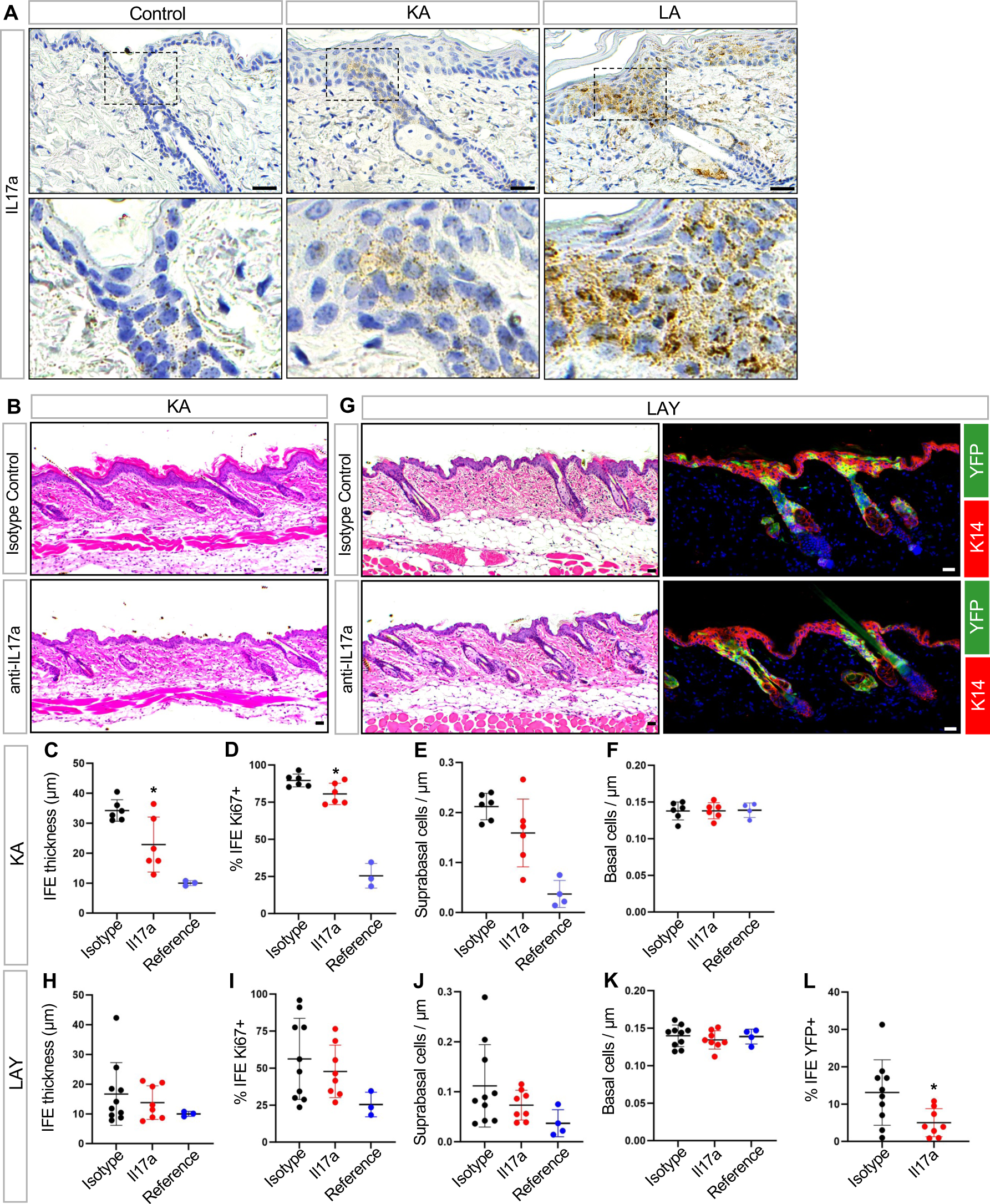
Il17a neutralization partially alleviates some disease phenotypes. **A.** RNA-scope *in situ* staining for *Il17a* in control, KA and LA skin, 2 weeks post-TAM. Bottom panels are magnified views. **B.** Histology of KA skin treated with anti-Il17a neutralizing antibody (bottom) or isotype control (top), 2 weeks post-TAM. **C.** Quantitation of IFE thickness in KA mice treated with anti-Il17a neutralizing antibody (red) or isotype control (black), 2 weeks post-TAM. Measurements from untreated control mice (blue) are shown as a reference. **D.** Same as (C), but with quantitation for basal IFE proliferation. **E.** Same as (C), but with quantitation for suprabasal IFE cell abundance. **F.** Same as (C), but with quantitation for basal IFE cell abundance. **G.** Histology of LAY skin treated with anti-Il17a neutralizing antibody (bottom) or isotype control (top), 2 weeks post-TAM (left panels). IHC for YFP+ mutant cells (green) in these samples (right panels). **H.** Quantitation of IFE thickness in LAY mice treated with anti-Il17a neutralizing antibody (red) or isotype control (black), 2 weeks post-TAM. Measurements from untreated control mice (blue) are shown as a reference. **I.** Same as (H), but with quantitation for basal IFE proliferation. **J.** Same as (H), but with quantitation for suprabasal IFE cell abundance. **K.** Same as (H), but with quantitation for basal IFE cell abundance. **L.** Same as (H), but with quantitation for basal YFP+ mutant cells in the IFE. *, p < 0.05 by unpaired t-test, comparing only samples from antibody-treated mice. For KA studies, n = 6 mice per treatment group. For LAY studies, n ≥ 8 mice per treatment group. Note that identical reference data are shown for C-F and H-K, respectively, and that reference data were not used for statistical comparisons. n = 3-4 mice for reference data. Scale bar, 50 μm.

To test the functional role of Il17a in disease, we treated KA mutant mice with neutralizing antibodies against this cytokine. Relative to isotype control, Il17a neutralization partially reduced IFE thickness and proliferation (**Figure 6B-D**). Reduced epidermal thickness was associated with fewer differentiated suprabasal IFE cells, without changes in the abundance of underlying basal progenitors (**Figure 6E-F**).

We next attempted to extend these findings to LAY mutant mice. However, unlike KA mice, we observed no significant effect of Il17a neutralization on IFE thickness, proliferation or suprabasal cell counts, features that are increased non-cell autonomously in LAY mutants (**Figure 6G-K**). Nonetheless, anti-Il17a treatment significantly inhibited the expansion of YFP-labeled mutant cells from the uHF into the IFE (**Figure 6L**). Together, these findings further demonstrate that targeted barrier disruption in either the IFE or the uHF causes complementary, but also distinct, skin phenotypes that are likely driven by non-overlapping disease mediators.

### Selective barrier disruption in K79+ uHF cells causes ichthyosis-associated phenotypes

As a final question, we circled back to our original observation that the uHF is lined by K79+ suprabasal cells, and asked whether even more precise ablation of *Abca12* in only these differentiated cells—the exact cells that express Abca12 in the uHF (**Figure 1D**)—can cause ichthyotic phenotypes. Since Lrig1-Cre^ERT^^2^ induces genetic recombination in stem cells that maintain the entire uHF, we therefore turned instead to mice expressing constitutive *Krt79* promoter-driven Cre (*K79-Cre* mice), which we previously generated and re-validated here (**Figure 7A**)^47^.

**Figure 7.**
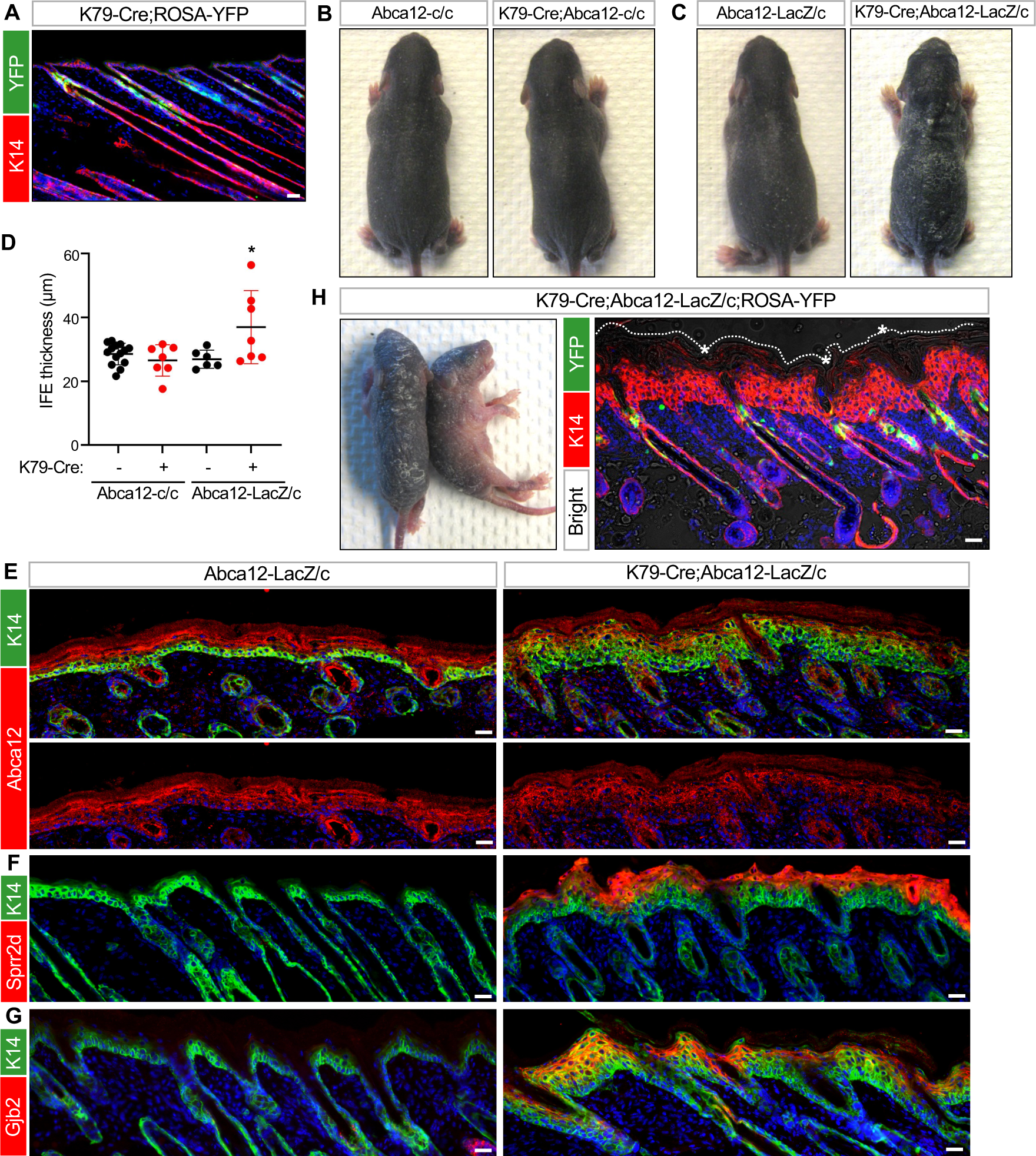
Targeted deletion of *Abca12* in K79+ uHF cells induces ichthyosis-associated phenotypes. **A.** IHC for YFP (green), confirming recombination in K79+ uHF cells in *K79-Cre;ROSA-YFP* mice at P12. **B-C.** Gross photos of P7 pups of the indicated genotypes. Only *K79-Cre;Abca12-LacZ/c* pups exhibited severe skin thickening and flaking. **D.** Quantitation of epidermal thickness for P7 pups of the indicated genotypes. Each data-point represents a single mouse. **E.** IHC for Abca12 (red), confirming that the protein was not ablated from the IFE of *K79-Cre;Abca12-LacZ/c* skin at P7. Bottom panels are single-channel views of Abca12 staining. **F.** IHC for Sprr2d (red) in *K79-Cre;Abca12-LacZ/c* or control skin at P7. **G.** Same as (F), but with staining for Gjb2 (red). **H.** Left panel, gross photos of two P6 mutant pups, both of genotype *K79-Cre;Abca12-LacZ/c* with *ROSA-YFP*. Right panel, IHC showing YFP expression (green) in developing hair canals after Cre-mediated recombination and thickened epidermis (red). Asterisk, hyperkeratotic material obstructing hair canals, visible by brightfield overlay. Dotted line denotes the top of the epidermis. See **Figure S4E** for IHC of littermate control skin. For **D**: *, p < 0.05 by one-way ANOVA with post hoc Tukey test, comparing *K79-Cre;Abca12-LacZ/c* pups against *Abca12-LacZ/c* or *K79-Cre;Abca12-c/c* pups. n ≥ 6 mice per genotype. Scale bar, 50 μm. See also **Figure S4**.

When *K79-Cre* was coupled with homozygous *Abca12* cKO alleles, some mutant pups exhibited very subtle skin flaking at P4-P7, which disappeared within days (**Figure 7B, S4A**). Since this mild phenotype may be due to inefficient Cre-mediated recombination (**Figure S4A**), we generated K79-Cre mice harboring both the *Abca12* KO (LacZ) and cKO (c) alleles, where deletion of only a single copy of *Abca12* is needed to fully ablate its expression (*K79-Cre;Abca12-LacZ/c* mice). In a subset of these mutant pups, we now observed more severe skin thickening and flakiness between P4-P7 (**Figure 7C-D**). We should note that some mutants did not exhibit overt phenotypes, while two mutants with thickened skin died just before P7 and were not included in our analysis. Finally, mutant animals that survived into adulthood appeared normal, suggesting that Cre ineffiency and variations in genetic background may have continued to cause incomplete penetrance (**Figure S4B-D**).

Focusing only on *K79-Cre;Abca12-LacZ/c* pups that exhibited ichthyotic features at P7, we confirmed that Abca12 was not deleted from the IFE, as expected (**Figure 7E**). Nonetheless, these pups possessed hyperplastic epidermis that upregulated Sprr2d, S100a9 and Gjb2, similar to LAY mutants (**Figure 7F-G**, **S3F**). As a final validation, we generated *K79-Cre;Abca12-LacZ/c* pups incorporating an additional *YFP* reporter allele. In these mutant pups, we again observed thickened epidermis and follicular hyperkeratosis. Critically, Cre-mediated recombination was highly restricted to developing hair canals, confirming K79-Cre targeting specificity (**Figure 7H**, **S4E**). In total, these findings firmly establish that K79+ cells in the uHF form a functioning barrier, and that highly targeted barrier disruption in this domain is sufficient to induce widespread ichthyosis-associated phenotypes.

## Discussion

Most studies on skin barrier function have focused on interfollicular epidermis, with less consideration for whether hair follicles play a role. Nonetheless, studies in human HI fetuses have reported that hyperkeratosis initially appears prior to barrier formation and is most pronounced in developing hair canals^48^. Concordantly, studies in *ichq* mice, which develop ichthyotic features due to a mutation in *Cst6*, have also found that disease onset coincides with the emergence of hair^49,50^. These and other findings support the notion that the formation of the hair canal—an event characterized by extensive cellular remodeling, as well as shifts in immune and microbial composition—represents a seminal but also potentially vulnerable event in skin development^19,51,52^.

Our initial observation that K79 is only expressed in the uHF, but not in the IFE, indicated that these domains are molecularly distinct^27,47^. Our results now argue that these domains are also functionally distinct. Previously, pharmacological studies have attempted to measure intrafollicular drug delivery, using varied approaches such as comparing haired versus non-haired skin, selectively blocking hair orifices, differential tape stripping and imaging^15,24^. These studies have suggested that hair follicles provide routes for drug penetration and may even act as drug reservoirs^23^.

Using a targetable genetic tool to disrupt lamellar granules and perturb barrier function, here we deduced at least 4 ways by which hair follicles modulate the skin’s barrier. First, we found that the uHF forms a functional, albeit “leakier,” barrier. Second, deleting *Abca12* in the uHF causes non-cell autonomous upregulation of barrier proteins in the IFE. Third, the uHF provides a passageway for sebum release, a process that requires constant desquamation in the hair canal. Fourth, uHF-derived mutant cells spread into the IFE. This last finding is contrasted by the observation that mutant cells in dorsal skin IFE are lost over time, leading to disease recovery. Differences in uHF versus IFE stem cell behavior, including proliferative potential, may account for these divergent outcomes^42,53,54^.

Additionally, we observed that the uHF is a site of enriched *Il17a* expression. Il17a is a critical proinflammatory cytokine implicated in numerous skin disorders including psoriasis, atopic dermatitis and ichthyosis^55,56^. In addition to Il17a, previous studies have reported that uHF keratinocytes also secrete factors such as Il7, Il15, Ccl2 and Ccl20 to recruit leukocytes^18,19,57^. Both T cells and Langerhans cells can be found in proximity to the uHF, further highlighting this domain as an important immune signaling center in the skin^18,57^.

Although HI is a skin-wide disease, our use of *Abca12* cKO mice helps untangle the contributions of IFE and uHF keratinocytes to barrier function. Our findings also suggest that perturbing one epithelial compartment in the skin can elicit compensatory responses from the other. Indeed, previous studies have reported that barrier dysregulation can cause upregulation of barrier-associated genes^58^, including *Abca12*^59,60^, and stimulate processes associated with cornification, such as lipid synthesis and LB secretion^5,32,61^. In keratinocytes, ceramide accumulation can act via PPAR nuclear receptors to upregulate *Abca12* and promote differentiation^62–64^. While these compensatory responses are typically regarded as cell autonomous phenomena, a non-cell autonomous effect acting broadly across different epithelial compartments, as seen here, has not been previously demonstrated.

Follicular hyperkeratosis and obstructed hair canals have been observed in several mouse mutants and human dermatoses. These include mice with mutations in genes encoding proteolytic enzymes such as matriptase, cathepsin L and matrix metalloproteinase 9, suggesting that these proteases are critical for desquamation^65–67^. Loss of phospholipase C and overexpression of Nrf2 have also been reported to induce ichthyotic phenotypes and occluded hair canals^68,69^. In patients, follicular hyperkeratinization is seen in disorders such as acne, keratosis pilaris, ichthyosis follicularis with atrichia and photophobia (IFAP), keratosis follicularis spinulosa decalvans and pityriasis rubra pilaris^70–72^. Follicular occlusion is also a unifying characteristic of a disease “tetrad” that includes hidradenitis suppurativa, acne conglobata, dissecting cellulitis and pilonidal sinus. In our studies, we found that targeted deletion of *Abca12* in the uHF, but not in the IFE, is sufficient to cause hair plugging. These effects are likely due, in part, to disrupted LG-mediated delivery of Klk6/7 and Ctsd, which are both normally enriched in K79+ cells of the uHF and serve to degrade corneodesmosomes^7,73^. Subsequently, sebum becomes trapped within the hair canal, where these lipids are unable to fulfill their moisturizing and anti-microbial functions^74–76^.

Recent gene expression studies across multiple ichthyosis subtypes have indicated that these disorders share molecular similarities with psoriasis vulgaris, including upregulation of IL17, IL36 and TNFα signaling pathways^44,77^. Moreover, disease severity correlated with IL17A serum levels and IL17 pathway gene expression^77^. However, a recent clinical study reported that secukinumab, a monoclonal antibody against IL17A, was ineffective at treating ichthyosis patients^78^. We also observed in mice that neutralizing Il17a yielded only partial skin improvement, suggesting that targeting multiple inflammatory pathways may be critical^43,79^. Finally, use of a NOS2 inhibitor or the JAK inhibitor tofacitinib has recently been shown to promote differentiation and barrier function in a 3D model of HI^44^. These findings raise the possibility that drugs which restore barrier function may one day provide a novel treatment strategy for HI and other ichthyosis subsets.

### Limitations of the Study

Ichthyoses are inherited diseases that affect the entire skin. Because our studies seek to identify compartment-specific contributions to the skin’s barrier, the KA and LA mice used in our experiments are chimeric for *Abca12* deletion and may not fully recapitulate all disease features. At the same time, the K14-CreERT and Lrig1-CreERT2 drivers that target *Abca12* deletion to the adult IFE and uHF, respectively, can induce occasional recombination outside of their expected domains. Thus, we cannot formally rule out the possibility that these occasional recombination events may contribute to disease phenotypes. In KAY mice, our analyses of hairless volar skin suggest that losing *Abca12* does not reduce the overall fitness of basal layer epidermal stem cells; nonetheless, we cannot rule out the possibility that Abca12 possesses vital skin site-specific functions, possibly explaining why mutant cells become lost from dorsal skin over time. Also currently unclear is why our use of the K79-Cre driver to delete *Abca12* causes incomplete disease penetrance. Finally, Il17a neutralization yielded modest effects, especially in LAY mice. This may be partially due to phenotypic variability in this strain, although other disease mediators besides Il17a are likely also involved and remain to be identified.

## ACKNOWLEDGEMENTS

We are grateful to the Dlugosz lab (University of Michigan) for insightful discussions, and to Debra Crumrine, Dr. Jason Meyer (Vanderbilt), and Sasha Meshinchi (University of Michigan) for TEM advice. We also thank Drs. Michael Fitzgerald and Mason Freeman (Harvard Medical School) for Abca12 antibody, and Thomas Huyge (University of Michigan) for clerical assistance. S.Y.W. acknowledges support from the NIH (R01AR080654 and R01AR065409), the LEO Foundation (LF18017), and the UM Skin Biology and Disease Resource-based Center (P30AR075043).

## AUTHOR CONTRIBUTIONS

Conceptualization and methodology, N.C.F., R.E.B., Y.U., S.Y.W. Investigation, N.C.F., R.E.B., M.T.M., J.K.P., A.L.M., N.A.V., D.J.G., S.Y.T., S.Y.W. Formal Analysis, N.C.F., R.E.B., M.T.M., J.K.P., S.Y.T., S.Y.W. Writing – Original Draft, Review & Editing, N.C.F., S.Y.W. Funding Acquisition and Supervision, S.Y.W.

## DATA AVAILABILITY

RNA-seq data generated for this study can be accessed through GEO: accession # GSE254889 (token: onqvqwqohdmdlcd).

## DECLARATION OF INTERESTS

The authors declare no competing interests.

Artificial intelligence (A.I.) was not used for the preparation of this manuscript.

## SUPPLEMENTAL FIGURE LEGENDS

**S1. Differentiation- and barrier-associated proteins are present in the uHF**

**A.** Co-localization of Klk7, Ctsd and K10 (red) with K79 (green) in the uHF. Lower panels are single channel views.

**B.** TEM of the IFE, with the right magnified panel showing secreted lamellar material (outlined in yellow) at the granular layer apical side. Scale bar, 200 nm.

**C.** TEM of hair follicle epithelium. Asterisk, hair shaft. Red, presumably K79+ granular cell abutting the hair shaft. Right and bottom magnified panels show lamellar material (outlined in yellow) being secreted into the interstitial space between the granular and cornified layer.

**D.** TEM of IFE-uHF interface, with magnified views of the IFE (top right) and hair follicle (bottom right). Brown line, IFE cornified layers. Green lines, hair follicle cornified layers. Note that the follicular cornified layers become thinner at the proximal end (bottom).

Scale bar for A, 50 μm. Scale bar for B, 200 nm. Scale bar for C, D, 1 μm.

**S2. Mouse models of *Abca12* deletion**

**A.** Schematic of *Abca12* KO allele containing a *LacZ* insertion between exons 3 and 4 (tm1a). Following FLP-mediated recombination, the KO allele is converted into a cKO allele with *loxP* sites flanking exon 4 (tm1c). After Cre-mediated recombination, exon 4 is removed, resulting in a non-functional gene (tm1d).

**B.** Genotyping strategy to identify wild-type (+), KO (tm1a) and condition (tm1c) alleles of *Abca12*. Please see methods section for primer sequences and PCR conditions.

**C.** IHC staining for Abca12 (red, left panels) and LacZ staining (blue, right panels) in control and *Abca12* KO skin. Arrow, *Abca12* promoter-driven LacZ expression in the uHF of developing follicles.

**D.** IHC staining for Abca12 (red) in control and *K5;Abca12-cKO* skin.

**E.** Gross images of nude mice grafted with control or *Shh;cKO* skin (left). Right, *Shh;cKO* graft, with IHC staining to detect *Shh* promoter-driven GFP (green, asterisk), identifying follicles of donor origin^47^. Arrow, hyperkeratotic uHF impeding hair shaft emergence, visible by brightfield overlay.

Scale bar, 50 μm.

**S3. Targeting *Abca12* deletion to different skin compartments and upregulation of S100a9**

**A.** Mice expressing K14-Cre^ERT^ and a YFP reporter (K14;YFP) exhibit recombination primarily in the IFE (left, green staining), whereas mice expressing Lrig1-Cre^ERT2^ and YFP reporter (Lrig1;YFP) exhibit recombination primarily in the uHF (right).

**B.** Gross image of shaved dorsal skin from an LA littermate control mouse for **Figure 2H**, 10 weeks post-TAM.

**C.** IHC for S100a9 in *Lrig1-Cre^ERT^*^2^*;Abca12-c/c;ROSA-YFP* (LAY) mice (bottom). Control mice are similar to LAY mutants, but possess one wild-type copy of *Abca12* (*Abca12-c/+*) (top). S100a9 upregulation (red) in the IFE of LAY skin does not overlap with YFP+ uHF cells (green).

**D.** IHC for YFP (green) and Abca12 (red) in KAY and control mice, 2 weeks post-TAM. Arrows indicate labeled control cells expressing Abca12 (top panels) or labeled mutant cells lacking Abca12 (bottom panels). A corresponding single-channel view of Abca12 staining is shown for each image.

**E.** Quantitation of recombined YFP+ cells in the uHF of LAY (red) or control (black) mice, showing that mutant cells persist in the uHF, up to 20 weeks post-TAM. n ≥ 4 mice per genotype, per timepoint.

**F.** IHC showing upregulated S100a9 (red) in the IFE of mutant *K79-Cre;Abca12-LacZ/c* skin at P7 (bottom). Littermate control skin lacks *Cre* (top).

Scale bar, 50 μm.

**S4. Adult *K79-Cre;Abca12-LacZ/c* mice do not exhibit overt skin defects**

**A.** Top, gross photos showing subtle skin flaking in a *K79-Cre;Abca12-c/c* mutant pup at P4 (right panel), with a littermate pup lacking Cre as a control (left panel). Bottom, IHC showing strong Abca12 expression (red) that overlaps with K79 (green) in the developing uHF (dotted lines) in control skin at P4 (left panels). Mutant skin at P4 exhibits heterogeneous Abca12 staining in the uHF (right panels). A corresponding single-channel view of Abca12 staining is shown for each image.

**B.** Skin histology from a 9 week old *K79-Cre;Abca12-LacZ/c* mouse (bottom) and control littermate lacking Cre (top).

**C.** Quantitation of IFE thickness in mutant mice (red) or controls (black) at 9 weeks of age. n = 5 mice per genotype.

**D.** Same as (C), but with quantitation for basal IFE proliferation.

**E.** IHC staining of P6 *Abca12-LacZ/c*;*ROSA-YFP* pup lacking Cre. This is a littermate control for animals depicted in **Figure 7H**. Scale bar, 50 μm.

## KEY RESOURCES TABLE

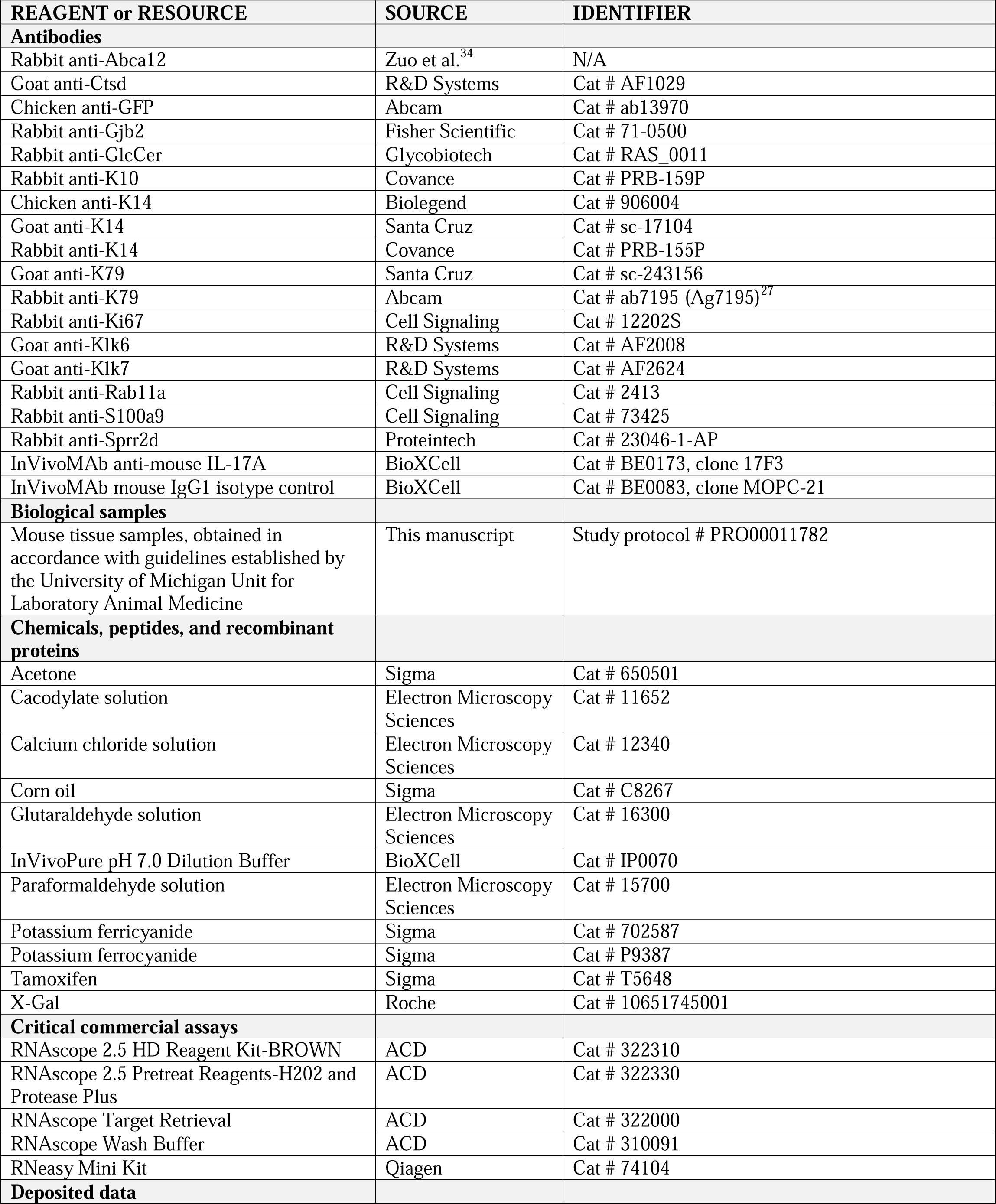
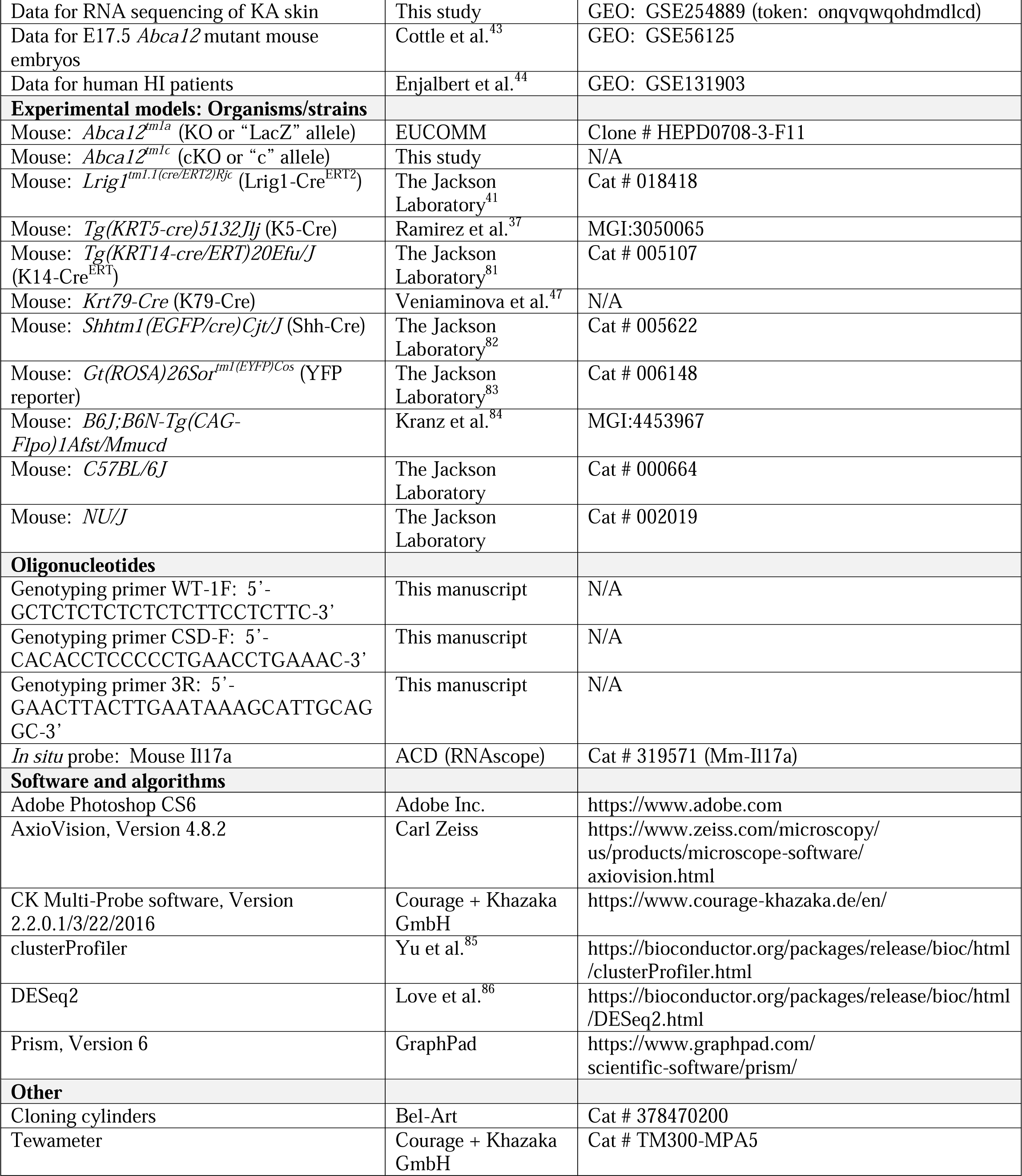

## STAR METHODS

## RESOURCE AVAILABILITY

### Lead contact

Further information and requests for resources and reagents should be directed to and will be fulfilled by the lead contact, Sunny Wong.

### Materials availability

All reagents generated in this study are available from the lead contact.

### Data and code availability

● RNA sequencing data generated for this study have been deposited in the GEO database: https://www.ncbi.nlm.nih.gov/geo/query/acc.cgi?acc=GSE254889 (token: onqvqwqohdmdlcd)
● This paper does not report original code.
● Any additional information required to reanalyze the data reported in this work is available from the lead contact upon request.

## EXPERIMENTAL MODEL AND STUDY PARTICIPANT DETAILS

### Animals

To generate *Abca12* KO (tm1a) mice, embryonic stem cells from clone HEPD0708-3-F11 were purchased from EuComm and microinjected into blastocysts. *Abca12^tm1a^* mice were crossed with Flpo mice (*B6J;B6N-Tg(CAG-Flpo)1Afst/Mmucd*) to generate *Abca12* cKO (tm1c) mice. To induce *Abca12* deletion in adults, 8 week old mice were injected intraperitoneally with tamoxifen dissolved in corn oil, at a dose of 1 mg/40 grams body weight for 3 consecutive days, and harvested between 2-20 weeks after the initial day of tamoxifen administration. For Il17a neutralization studies, mice were injected intraperitoneally with 250 μg of anti-Il17a antibody or IgG1 isotype control, diluted in 200 μL InVivoPure dilution buffer, once every 3 days starting 1 day after the final dose of tamoxifen, and then harvested 2 weeks after the initial day of tamoxifen administration. For grafting experiments, dorsal skin from *Shh;Abca12-cKO* or littermate control pups was removed and sutured onto the backs of female athymic *NU/J* mice. All *Abca12* mutant mice were of a mixed genetic background, and both genders were analyzed in roughly equal numbers for experiments. *C57BL/6* mice of both genders were analyzed by immunohistochemistry and transmission electron microscopy at 8 weeks of age, and for barrier function at the ages indicated in the text. All mice were used in accordance with regulations established by the University of Michigan Unit for Laboratory Animal Medicine.

## METHOD DETAILS

### Immunohistochemistry and RNA *in situ* staining

Frozen sections were stained by IHC using the following antibodies: goat anti-K79 (1:300); rabbit anti-K79/Ag-7195^27^ (1:500); goat anti-K14 (1:1,000); chicken anti-K14 (1:1,000); rabbit anti-K14 (1:1,000); rabbit anti-Abca12 (1:2,000)^34^; rabbit anti-Rab11a (1:100); goat anti-Klk6 (1:100); goat anti-Klk7 (1:100); goat anti-Ctsd (1:100); rabbit anti-GlcCer (1:1,000); rabbit anti-K10 (1:1,000); rabbit anti-Sprr2d (1:200); rabbit anti-Gjb2 (1:2,000); rabbit anti-S100a9 (1:500); chicken anti-GFP/YFP (1:1,000); and rabbit anti-Ki67 (1:500). Multiple IHC images from the same field were captured and merged using the Auto-Blend feature of Adobe Photoshop CS6 to maximize image sharpness automatically across multiple focal planes. For RNA *in situ* staining, 5 μm paraffin skin samples were stained with a probe against mouse *Il17a* (Mm-Il17a) using RNAscope 2.5 Brown kit, according to manufacture’s instructions.

### Transmission electron microscopy

For TEM, skin was fixed in 2% paraformaldehyde, 2.5% glutaraldehyde, 0.1 M cacodylate and 2 mM calcium chloride for 2 hours at room temperature. The fixative was replaced, and samples were incubated overnight at 4 degrees C. The next day, samples were rinsed and stored in 0.1 M cacodylate buffer at 4 degrees C. Samples were processed for TEM using standard protocols, sectioned at 70 nm thickness, and imaged using a JEOL JEM 1400 microscope at an HT voltage of 60 kV. Images were collected using AMT Capture software.

### Barrier Assays

Newborn pups were decapitated and submerged below the neck in 1 mg/ml X-gal diluted in 5 mM potassium ferrocyanide and 5 mM potassium ferricyanide, pH 4.5, overnight at 37 degrees C. To assess barrier function in adults, mice were euthanized and immediately shaved. A ∼2 x 2 cm region of telogen skin was swabbed with a Q-tip (paper cotton swab) repeatedly dipped in 100% acetone for 5 minutes. Both treated and untreated skin were then excised and flattened against a paper towel. A cloning cylinder was affixed to the skin surface with lubricant (white petrolatum), the chamber was filled with 200 μL X-gal solution, and samples were floated on PBS inside a 6-well plate and incubated overnight at 37 degrees C. To visualize samples by whole-mount, cloning cylinders were removed the next day, and the samples were rinsed briefly in PBS. Skins were re-stretched on a paper towel, patted dry, covered with Elmer’s rubber cement, allowed to dry for 5 minutes, and covered with transparent tape. The samples were floated epidermis-side up for 6 hours in 5 mM EDTA/PBS at 37 degrees C, and the epidermis was peeled away from the dermis. Separated epidermis was rinsed in PBS, fixed in formalin for 30 minutes, rinsed again, and imaged. For TEWL, mice were anesthetized, dorsal skin was shaved, and loose hairs were gently removed with a Kimwipe. Mice were allowed to acclimate in an upright resting position for 5 minutes, and telogen skin was probed using a Tewameter TM300-MPA5 and CK Multi-Probe software. Littermate mice lacking *Cre* or possessing one wild-type copy of *Abca12* were designated as controls.

### RNA extraction and sequencing

RNA was harvested from 4 KA mice and 3 littermate controls, 2 weeks post-TAM. The epidermis was isolated from telogen skin, as previously described^39^, and processed using RNeasy mini kit. RNA-seq was performed by Novogene Corporation, and data were processed using DESeq2^86^ to identify differentially expressed genes (DEGs) with |log2 fold-change (FC)| > 1 and adjusted p-value < 0.05. R package clusterProfiler^85^ was used to test statistical enrichment of DEGs in KEGG pathways. For overlap analysis, we used data from HI patients generated by Enjalbert et al.^44^ (GSE131903), |log2FC| > 1 and adjusted p-value < 0.01; and data from E17.5 *Abca12*-mutant mouse embryos generated by Cottle et al.^43^ (GSE56125), FC > 1.6 and adjusted p-value < 0.05. Analyses were performed using GEO2R. Venn diagrams were generated using http://bioinformatics.psb.ugent.be/webtools/Venn/

### Abca12 genotyping

PCR was performed on DNA isolated from ear tissue. For tm1a and tm1c: WT-1F (5’-GCTCTCTCTCTCTCTTCCTCTTC-3’), CSD-F (5’-CACACCTCCCCCTGAACCTGAAAC-3’), and 3R (5’-GAACTTACTTGAATAAAGCATTGCAGGC-3’) primers were used together with an annealing temperature of 56° C and 40 amplification cycles. See also **Figure S2A-B**.

## QUANTIFICATION AND STATISTICAL ANALYSIS

### Quantitation

Frozen sections stained for Ki67 and YFP were quantitated for proliferation from 3 random fields at 20x magnification. Only YFP-negative, basal layer IFE cells were manually counted to assess non-cell autonomous effects. To quantitate basal and suprabasal cells, the number of DAPI+ nuclei located either at the lowest layer of the IFE, or above that layer, respectively, were counted and normalized by the length of the IFE. To quantitate labeled uHF cells that had entered the IFE in LAY mice, the number of YFP+ basal cells in the IFE was counted and normalized to total IFE basal cells from 3 random fields. To measure IFE thickness, the vertical length of the IFE was measured at 3 random locations per sample and averaged. For sebaceous gland quantitation, 10 representative H&E fields were imaged at 20x magnification, each gland was manually traced, and the total SG area was calculated in pixels using AxioVision software. For adult studies, only telogen skin samples were quantitated.

### Statistics

Unpaired student’s t-test was performed at the following website: http://www.physics.csbsju.edu/stats/Index.html. One-way ANOVA was performed at the following website: https://www.socscistatistics.com/tests/anova/default2.aspx. All data-points shown represent independent mice.

