## Supplementary figures and images for "Hair follicles modulate skin barrier function"

### Supplemental Figure S1

Ford, Benedeck_Fig S1

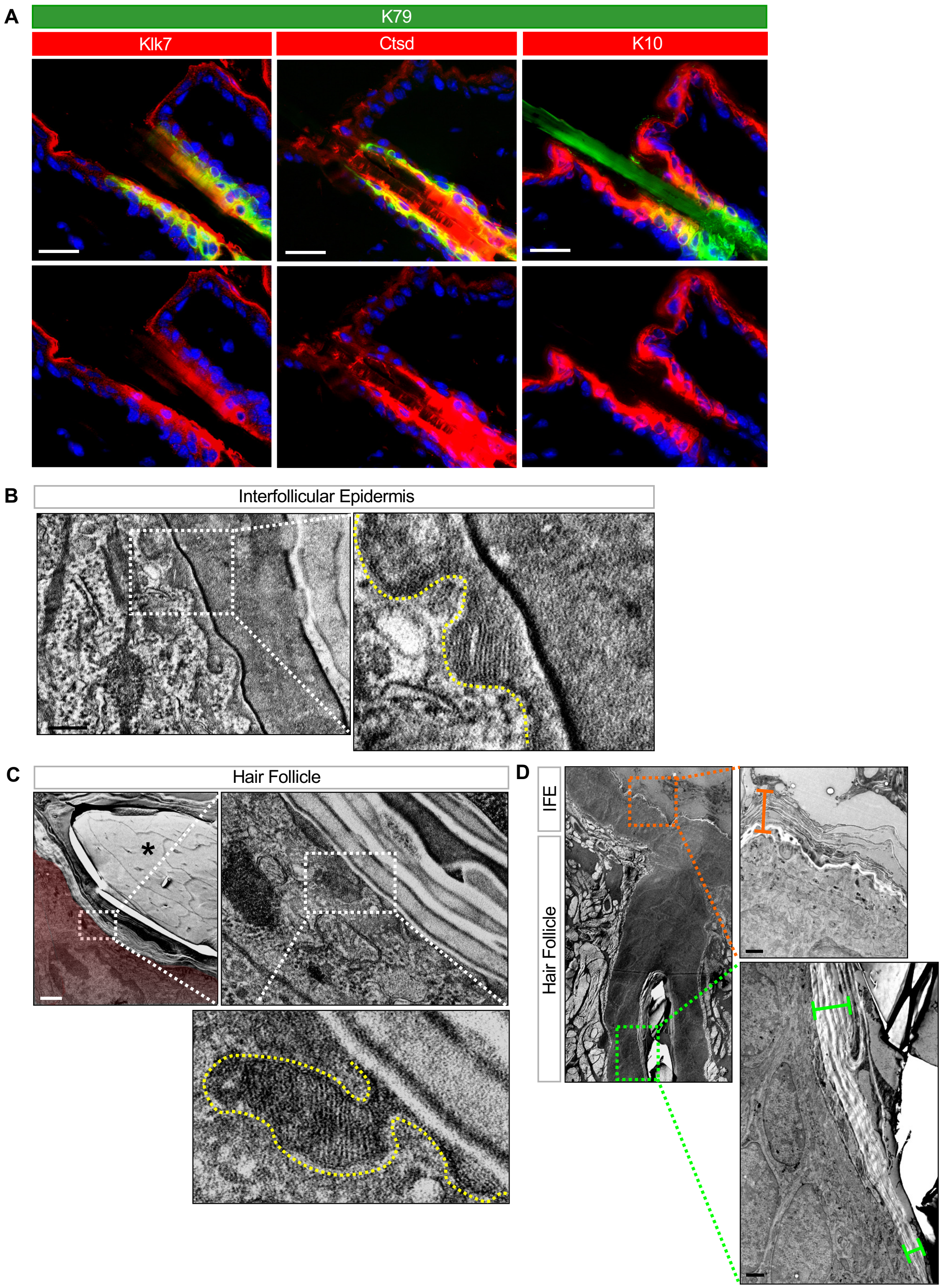

### Supplemental Figure S2

**Ford, Benedeck_Fig S2**

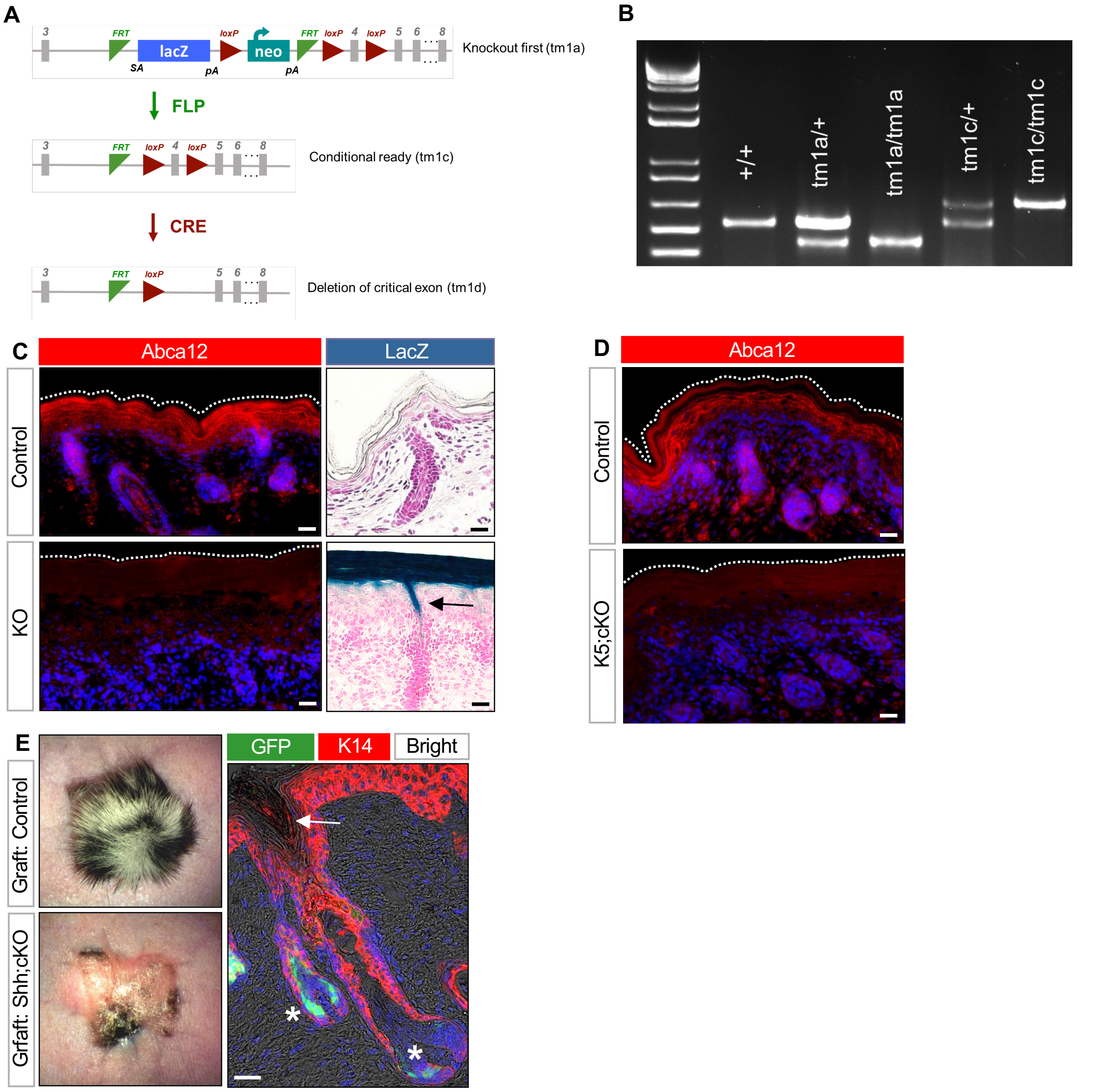

### Supplemental Figure S3

**Ford, Benedeck_Fig S3**

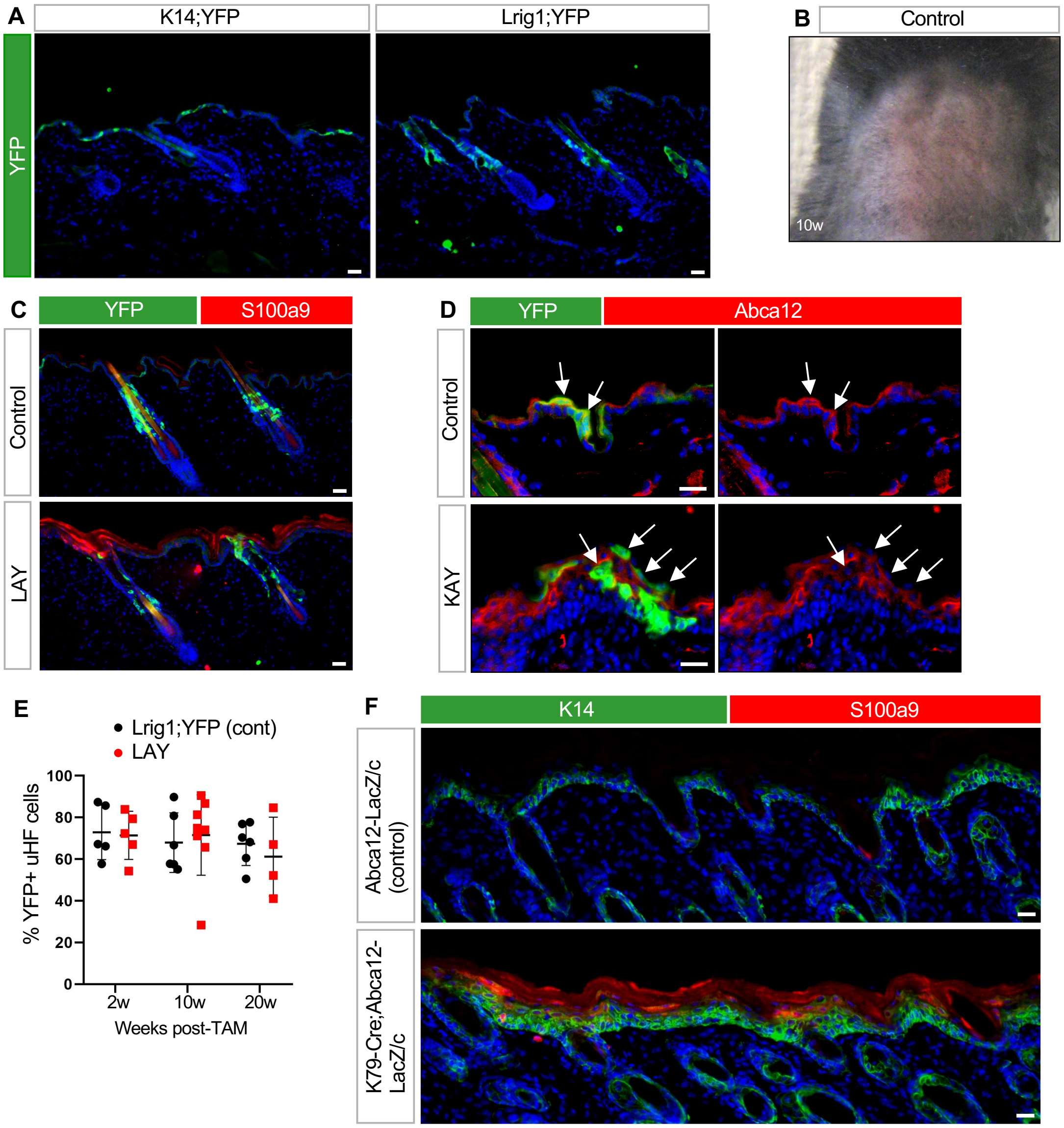

### Supplemental Figure S4

**Ford, Benedeck_Fig S4**

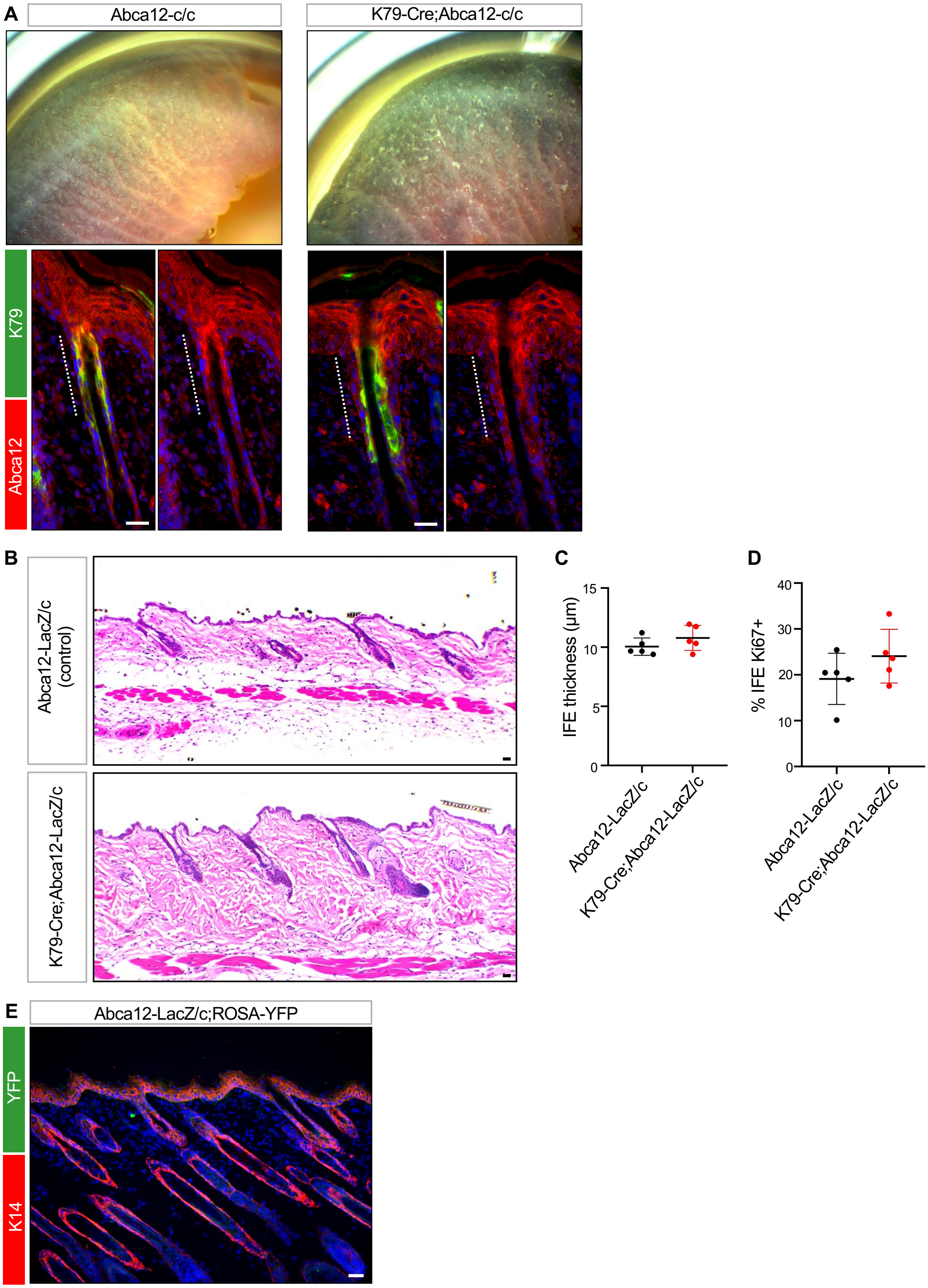
